# Apparent generalism in *Pseudomonas aeruginosa* is underpinned by Convergent, Cryptic Specialization

**DOI:** 10.64898/2026.05.15.725152

**Authors:** EC Mehlferber, I. Irby, M. Yarter, N. Lowery, R. Lowhorn, Y. Appaji, J. Eum, H. Song, B. Stone, SP Brown

## Abstract

Microbes that span environmental reservoirs and diverse human infections are often described as generalists or as ubiquitous, yet ecological theory predicts generalism should be unstable when specialists outperform within any one niche. Here we show that *Pseudomonas aeruginosa*’s apparent ecological breadth reflects convergent, cryptic specialism (CCS) – repeatable, environment-linked genomic differentiation that is not confined to deep lineages – rather than strict specialism or generalism. An analysis of 6,627 genomes with source-environment metadata reveals that environments are broadly dispersed across the phylogeny, inconsistent with lineage-locked specialization. Despite this shallow phylogenetic structure, genome content predicted environment-of-isolation across nine genotype-discernible environments, including under cross-validation that blocks phylogenetic relatedness. Interpretable feature profiles recovered both shared and environment-specific signals, including genomic signatures distinguishing distinct chronic and acute human infections. Finally, phenotypes from 47 diverse strains clustered more strongly by environmental source than by phylogenetic relatedness. Together, these results indicate that widely distributed bacterial “generalists” can comprise mixtures of cryptic, convergent specialists. By mapping genotype-defined ecological structure, our approach can identify when apparent generalists harbor hidden structure relevant to infection risk.

## Main

From humans to microbes, organisms face tradeoffs in ecological breadth^1–6^, choosing between doing a few tasks well (specialists) or many tasks tolerably (generalists). This tradeoff creates a longstanding paradox: if specialists outperform generalists in any given environment, why do generalists persist? Classic theory answers that generalism is favored when resources are patchy^3^ or conditions fluctuate over time^4^. Microbial evolution studies refine this picture by showing that when populations experience sustained selection in a single setting, ecological breadth can erode as lineages locally adapt and tradeoffs accumulate^7,8^. These results highlight a key inference problem: when a microbe is observed across diverse habitats (including humans), does this breadth reflect a genuinely broad niche, or the aggregation of multiple emerging niche specialisms, including pathogenesis? Motivated by these questions, we use rapidly expanding bacterial genomic resources to re-examine the generalism–specialism tradeoff, with unprecedented genomic and environmental resolution. We focus on *Pseudomonas aeruginosa* (PA), an environmental microbe and model opportunistic pathogen found in environmental and human-impacted sites, as well as clinical settings where it causes acute and chronic infections in diverse body sites, notably in the lungs of people with cystic fibrosis (CF).

We evaluate three hypotheses for the extent of specialism versus generalism in PA. First, strict specialism (Fig. 1ai) proposes that lineages acquire and lock in environment-specific genetic adaptations, leading to strong environmental clustering within lineages on the evolutionary tree^7^,^8^). Second, strict generalism (Fig. 1aii) posits that strains are functionally equivalent and interchangeable across environments. Niche breadth arises from plastic or broadly effective traits, leading to an absence of reproducible, phylogenetically independent genetic associations with environment. Finally, we propose a third model, convergent cryptic specialization (CCS; Fig. 1aiii), where environment-specific genetic filtering and adaptation is detectable (rejecting strict generalism), but also rapidly erased on phylogenetic timescale (rejecting strict specialism). CCS therefore predicts shallow phylogenetic structure alongside robust gene-environment associations.

A precedent for such genetic convergence exists in the CF lung, where strains from across the PA phylogeny have been reported to repeatedly evolve similar and diagnostic traits^9–13^. Here, we critically evaluate the CCS model across diverse sample environments. By distinguishing between strict specialism, strict generalism, and CCS, we aim to clarify how broad ecological breadth can be sustained within a single bacterial species.

### Environmental associations are phylogenetically shallow

Strict specialism (H1, Fig. 1Ai) predicts deeper phylogenetic structuring of environmental associations, with related strains more often occupying similar environments. To evaluate this prediction, we first asked whether environmental origin showed deep or shallow conservation across the largest clade of PA. We constructed a maximum likelihood phylogeny from 11,401 genomes downloaded and quality filtered from NCBI and annotated with environmental metadata (Fig. 1b). PA comprises two major clades, A and B, following the reclassification of Clade C as *P. paraeruginosa*. Clade B is dominated by a small number of phylogenetically distinct near-clonal lineages, whereas Clade A encompasses substantially greater diversity and the majority of sampled isolates (Supplementary Fig. 2). Accordingly, we restricted primary analyses to Clade A (see Supplementary Fig. 3 for limited model transfer to clade B). Visualization of the Clade A phylogeny with an environmental heatmap showed that environments of isolation are broadly distributed across the tree (Fig. 1b), in contrast to the lineage specialization prediction of deep phylogenetic structuring by environment (Fig. 1ai). We quantified phylogenetic clustering using trait depth (consenTRAIT τD^14^), estimated as the mean branch length from the root of clades in which at least 90% of members share a trait to their tips. Across all environments, phylogenetic depth was no deeper than expected under a null distribution generated by randomizing environment labels across the tree (and was significantly *less* deep for 4 environments), consistent with frequent transitions among environmental groups (groupings defined quantitatively later in the paper).

**Figure 1.**
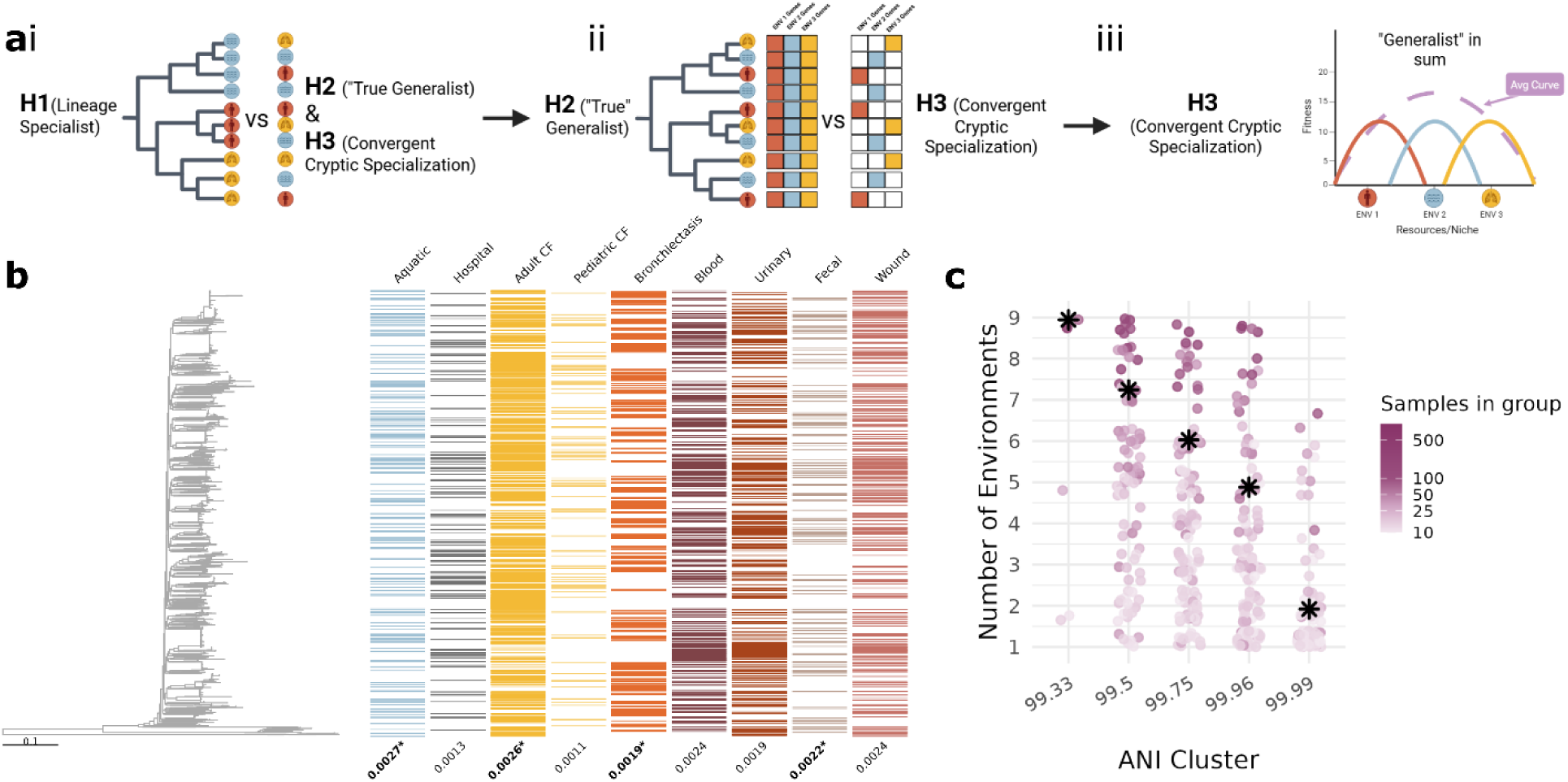
Environmental distributions show shallow phylogenetic structure, rejecting lineage specialization. (a) Schematic overview of three hypotheses for ecological specialization in P. aeruginosa. H1: Strict specialism (Fig. 1ai) predicts that lineages acquire and retain environment-specific adaptations, producing strong phylogenetic clustering of environments. H2: Strict generalism (Fig. 1aii) predicts that strains are functionally equivalent across environments, with no phylogenetic signal of environmental origin. H3: Convergent cryptic specialization (CCS) (Fig. 1aiii) proposes that strains independently adapt to similar selective pressures across the phylogeny, generating shared environment-linked signatures without deep phylogenetic structure. (b) Maximum-likelihood phylogeny of Clade A constructed from 6,627 genomes annotated with environmental metadata. The heatmap shows each strain’s environment of isolation. Trait depth (consenTRAIT τD, values below heatmap) was used to quantify phylogenetic associations. τD has the same units as the tree’s branch lengths (substitutions/site); smaller τD means shallower clustering (closer to tips) and weaker phylogenetic structure. Several environments, including Adult CF, Bronchiectasis, Fecal, and Aquatic, showed significantly lower trait depths than expected under randomization (bold, asterixed values). (c) Environment-of-isolation diversity across hierarchical ANI clusters 99.33%–99.99%, excluding clusters with <9 samples at each level (see Supplementary Fig. 1 for unfiltered counts). Even at the highest ANI thresholds, well-sampled clusters typically include strains from multiple environments, indicating limited ecological specificity at fine phylogenetic scales.

To examine finer-scale environmental distribution patterns that are not easily visible in Fig. 1b, we quantified environment-of-isolation diversity within closely related groups of strains (Fig 1c; grouping via average nucleotide identity (ANI) thresholds). At 99.3% ANI, nearly all groups spanned all environments; only a few small (poorly sampled) clusters were restricted, and even these occurred in at least two environments. At the genomovar level (99.5% ANI), and contrary to expectations of strong habitat restriction at this scale^15,16^, well-sampled genomovars contained strains from most or all environments (Fig. 1c; environmental membership for major genomovars in Supplementary Fig. 4). Increasing the ANI threshold did not reveal hidden specialization: even at 99.99% ANI, clusters typically included strains from at least two environments.

Overall, Figure 1 provides no evidence for deep phylogenetic conservation of environment, rejecting H1 and motivating tests of strict generalism (H2) versus convergent cryptic specialization (H3).

### Genomic content predicts environmental origin

Figures 1b, c provide evidence against strict specialism but remain consistent with both strict generalism (H2) and convergent cryptic specialization (CCS, H3), as both models predict that all lineages can occupy the full range of *P. aeruginosa* environments (Fig. 1a ii, iii). The key distinction is that CCS posits that genotype-environment associations can arise due to within-environment selection, environmental filtering or both (Fig. 1a iii).

We therefore used a supervised machine-learning classifier as a hypothesis test: can genomic features predict a strain’s environment of origin better than chance? This setup yields clear, falsifiable performance expectations: accuracy should be indistinguishable from a randomized baseline under H2, but above baseline under H3. To test our hypotheses, we trained an XGBoost^17^ classifier to predict each strain’s environment of isolation from the presence or absence of protein coding variants, grouped by 100% amino acid identity^18^.

We tested the CCS hypothesis across multiple environmental splits to determine which environmental separations are best supported, and in all cases we found support for the CCS hypothesis, with protein-coding variants predicting multiple source environments with accuracy in clear excess of randomized baseline (Fig. 2a, c, Supplementary Fig. 5). Fig. 2a summarizes our approach. Because phylogenetic structure was confined to a limited signal only below the genomovar level (Fig. 1c), we used genomovar-blocked cross-validation^19^ throughout our model training. We trained multiple rounds of XGBoost models in which each genomovar was withheld as a test set, which provided a conservative way to ensure that the model could not rely on any phylogenetic confounding or idiosyncratic cohort structure tied to particular genomovars.

**Figure 2.**
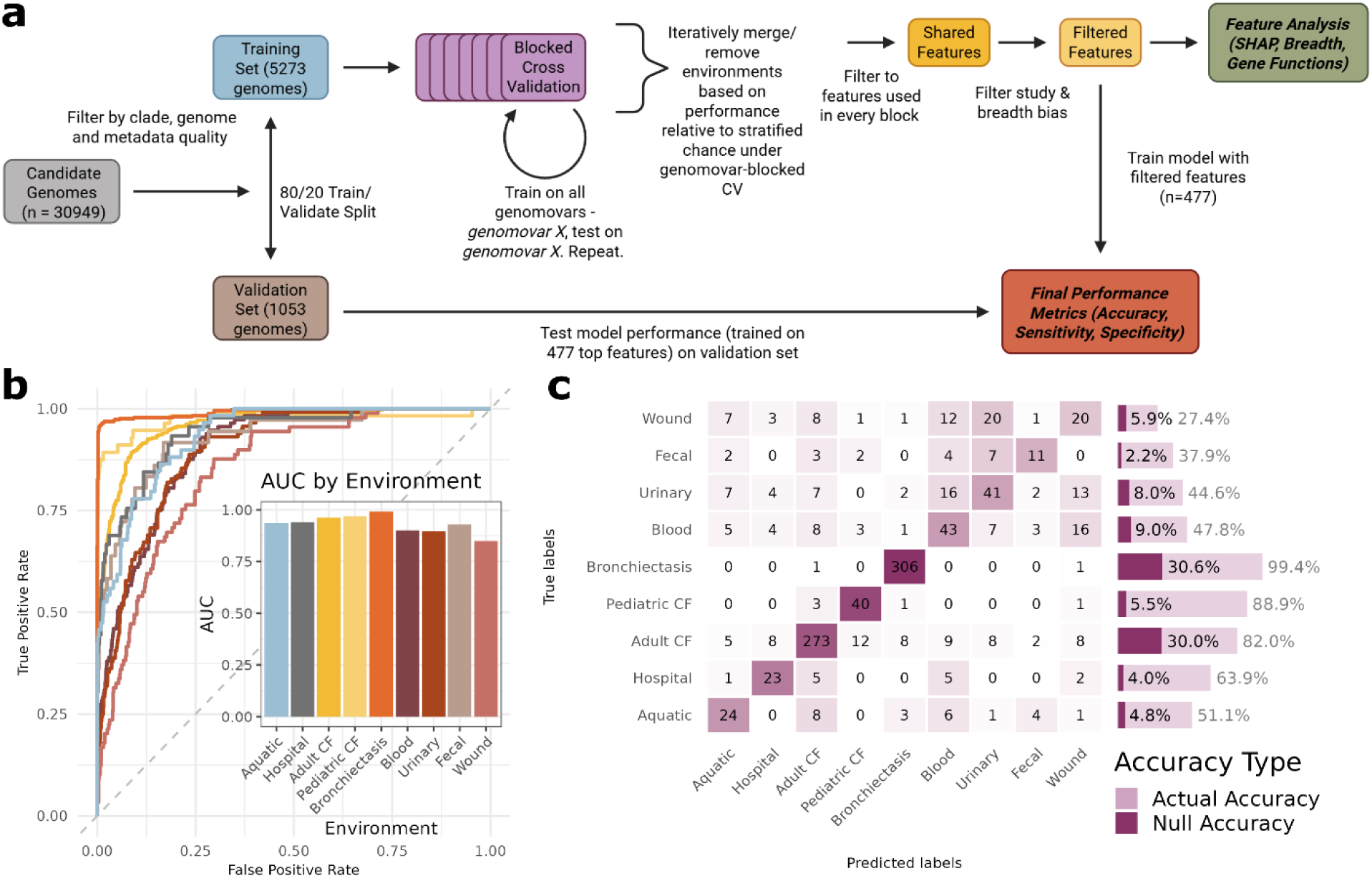
Environment of isolation is predictable from genomic content, supporting CCS. (a) Workflow diagram. Genomes were first filtered based on quality and relevant metadata, then split into training and final validation sets for XGBoost model training and evaluation. Blocked cross-validation was performed on the training data across hierarchical environmental groupings to identify which environments or clusters were best supported (Supplementary Fig. 5). Protein-coding variants were then filtered to retain those present or absent in >25% of genomovars (depending on if the model expected presence or absence for that environment) and to remove potential study-associated biases (for example, consistent sequencing or assembly errors within single studies). The resulting set of 477 protein-coding variants was then used to produce final predictions to be evaluated on held-out validation dataset (results in (b,c)). (b) Area under the receiver operator curve (AUC) plot showing validation model performance across True Positive and False Positive thresholds in a one versus rest comparison. The inset panel reports the AUC for each environment, demonstrating consistently high separation of each environment from the rest. (c) Confusion matrix showing validation model performance. Model accuracy per environment is shown to the right of the confusion matrix with comparison to expected performance under random assignment.

Starting from 17 metadata-defined categories, we treated environment definition as a discovery problem: which contrasts are reproducibly separable from genome content? Under genomovar-blocked CV, we computed normalized accuracy, defined as the observed accuracy minus the stratified-chance expectation (the accuracy a classifier would achieve by randomly guessing labels in proportion to their frequencies). Categories whose normalized accuracy hovered near zero therefore performed no better than chance. We retained only those categories that consistently exceeded this baseline across resamples. (Methods and Supplementary Fig. 5). This procedure yielded nine “genotype-discernible” environments, the set of environments that *P. aeruginosa* effectively “sees” as distinct given its genomic variation. The final set comprised three chronic lung environments (Adult CF, Pediatric CF, and non-CF bronchiectasis, henceforth termed Bronchiectasis), four likely acute non-lung human infection environments (Blood, Urinary, Fecal, and Wound), and two non-host environments (aquatic and hospital).

Prior to final testing of the nine-environment model on the fully held-out validation data, we applied several filtering steps to enrich for common and informative protein-coding variants. From our initial pool of 2773 protein-coding variants used across all blocks of the cross-validation, we first removed variants flagged as study-specific (potentially including cohort- or pipeline-specific sequencing or processing artifacts), defined as those showing >50% relative-abundance differences between the highest- and second-highest-abundance studies. Finally, we filtered to retain only variants whose predicted state was sufficiently common: variants predicted as present required presence in ≥25% of genomovars, and variants predicted as absent required absence in ≥25% of genomovars. After these steps, 477 protein-coding variants remained (Supplementary Table 1). Using this filtered set, we trained the final model and evaluated it against the fully withheld validation data (Fig. 2b, c).

Using this filtered feature set, the final model showed high one-vs-rest discrimination across environments (Fig. 2b, ROC AUC ∼.85–.99) indicating that each environment carries distinct, separable genomic signals, supporting our core CCS hypothesis. Our final validation analysis also demonstrates robust accuracy exceeding the null expectation (shuffled labels) across all environments (Fig. 2c). Performance in both AUC analyses and model accuracy is particularly strong for the chronic lung categories (Bronchiectasis, 99.2% accuracy, 0.992 AUC; Pediatric CF, 88.9% accuracy, 0.971 AUC; Adult CF, 82.0% accuracy, 0.962 AUC).

Acute infection categories exhibit higher misassignment rates among themselves, yielding lower accuracies (Blood, 47.8%; Urinary, 44.6%; Fecal, 37.9%; Wound, 27.4%) (Fig. 2c). This contrasts with their relatively high AUC values (Blood, 0.901; Urinary, 0.897; Fecal, 0.929; Wound, 0.849), indicating that while they are distinct from other environments, they share overlapping genomic signals that limit discrimination among acute infection classes. Sample environment assignment probability distributions (Supplementary Fig. 6) further support this pattern: acute categories show elevated within-group cross-prediction probabilities, consistent with overlapping selective pressures and/or smaller sample sizes for acute infection classes.

Across all environments, final validation model accuracy is 74.2%, with a weighted-F1 of 73.8%, a macro-F1 of 59.1%, and a balanced accuracy of 55.4% (Fig. 2C). Together, these results refute the prediction of genomic equivalence under generalism (H2) and support the CCS prediction that shared selective pressures produce reproducible genomic signatures consistent across phylogenetic backgrounds.

### Consistent mutational patterns are associated with environmental origin

Having established the predictive power of genomic signatures (Fig. 2b, c), we next examined the 477 filtered protein-coding variants driving this signal. To evaluate feature importance, we used SHAP scoring^20^. SHAP scores quantify how much each feature increases or decreases the model’s predicted probability for a given environment, relative to the model’s baseline prediction^20^. We computed SHAP values across all genomovar holdouts from the blocked cross-validation step to provide a comprehensive assessment of variant importance.

Figure 3A shows the most influential variants by global mean absolute SHAP (vertical lines) and their environment-specific importance (ΔSHAP per environment; colored dots). The histogram backdrop illustrates that low-importance coding variants collectively contribute more than any single top-ranked variant. Top features included wild-type (PAO1-matching) *mucA*, positively associated with acute infections, especially wounds, and negatively with chronic lung infections, particularly in Adult and Pediatric CF. This aligns with prior reports linking mucoidy (loss of function mutations in *mucA*) to chronic CF infections^21–23^ while extending the pattern to Pediatric CF and non-CF Bronchiectasis. Other key signals include mutant *tonB2* (iron and sulfate associated^24,25^; Pediatric CF-specific), wild-type *mexZ* and *mexB* (efflux regulation and antibiotic resistance^26,27^), mutant *fha1* (T6SS-associated^28^), and wild-type *lasR* (quorum sensing^29–31^).

**Figure 3.**
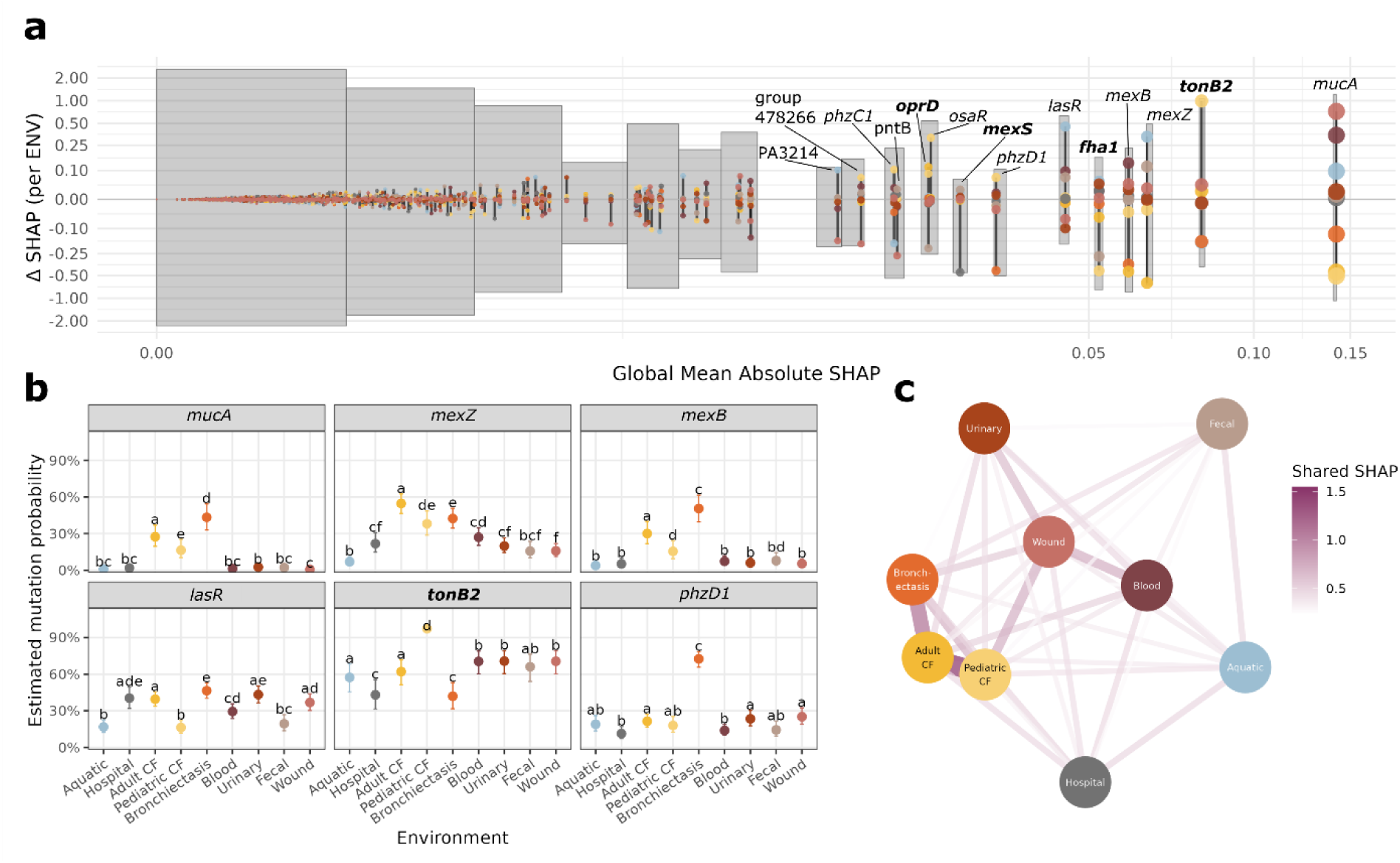
Model features reveal distinct and convergent genetic signatures of local environmental origin. (a) Top environment-discriminating genes, ranked by global mean absolute SHAP value (total importance across environments) and ΔSHAP (SHAP when variant is present minus SHAP when variant is absent within environments). This visualization highlights the most informative variants (*x-*axis, vertical lines) and their directionality for environmental classification (*y-*axis, circles). WT (PAO1) variants are labeled in regular text while specific mutants (ie. not WT relative to PAO1) are highlighted in bold. In the background is a histogram of total ΔSHAP across regions of the global mean absolute SHAP distribution, which provides context for the cumulative importance contributed by the many lower importance protein-coding variants. (b) Estimated mutation probability (probability that a gene differs from PAO1 WT) from the LMER mixed-effects model, shown as points (± SE) by environment across all *P. aeruginosa* genomes relative to the PAO1 reference (See Supplemental Figs. 7-8 for analysis restricted to putative Loss-of-Function mutations). The model includes *environment* as a fixed effect and *genomovar* as a random effect, with Tukey post hoc tests assessing pairwise differences. (c) Network visualization (force directed graph) of gene variants with consistent SHAP direction shared between environments, where edges represent the value of the shared SHAP score for each environment pair, computed by identifying all important genes with shared directions and summing the mean SHAP values for the predicted state (presence or absence) across those shared genes. This highlights that chronic lung infections cluster together and are more separated from acute infections.

To examine the biological basis of these SHAP patterns and confirm that they reflect functionally relevant genomic variation, we analyzed mutation profiles for the top predictive genes across environments (Fig. 3B, Supplementary Fig. 7). Using our mutation categorization pipeline^32^, we analyzed all 6,627 genomes relative to the PAO1 reference, classifying mutations as putative functional changes (amino acid substitutions) or loss-of-function variants (frameshifts or deletions). We then fit a LMER model with environment as a fixed effect and genomovar as a random effect to test for differences in mutation rates across environments, followed by Tukey post hoc comparisons (Fig. 3B, Supplementary Fig. 7).

Our mutation analysis shows that *mucA* mutations are significantly and substantially enriched not only in CF but across all chronic lung infection environments, consistent with our SHAP results (Fig. 3B), with further significant enrichment of putative loss-of-function mutations (Supplementary Figs. 7-8). We find a similar pattern of broad chronic-lung enrichment for mutations in both *mexZ* and *mexB*, which are significantly enriched in Adult CF and Bronchiectasis (Fig. 3B), again with additional enrichment of putative loss-of-function mutations (Supplementary Figs. 7-8).

Our SHAP analysis flags specific marker genes that enable reliable discrimination among chronic infections (notably *tonB2* and *phzD1*, Fig. 3a). Our mutation analysis (Fig. 3b) confirms that WT *tonB2* is significantly depleted in Pediatric CF relative to all other environments, while WT *phzD1* (involved in phenazine biosynthesis, ie. pyocyanin^33–35^) is significantly depleted in Bronchiectasis. Both genes exhibited frequent “contig-split” mutations in these environments, but these mutation patterns remain qualitatively consistent if contig splits are removed (Supplementary Fig. 9).

Finally, we flag that one of the most widely reported targets of CF mutations, *lasR* ^13,30^, is in fact broadly mutated across environments (Fig. 3b, Supplementary Figs. 7-8), extending recent findings (across more limited datasets) that disruption of QS regulation is not exclusive to CF-associated lineages^36,37^.

Our analyses in Fig. 3a,b indicate that certain environments have shared predictors (eg. chronic lung infections are linked by mutated *mucA*). To take a broader view on these links across all 477 protein-coding variants (including 118 hypothetical proteins in the top 10% of mean absolute SHAP per environment, and 283 in total, see Supplementary table 1), we summarized inter-environment relationships using a force-directed network (Fig. 3c).

Edges reflect summed SHAP values from directionally consistent protein-coding variants, calculated by matching variant prediction states (presence/absence) across pairs and averaging SHAP magnitudes. The resulting network shows clustering of chronic lung infections, Adult and Pediatric CF most closely, with non-CF Bronchiectasis adjacent.

Acute infections (Urinary, Wound, Blood) form a looser group, while Fecal, Hospital, and Aquatic environments are more distinct. The environmental organization in Fig. 3c potentially reveals *P. aeruginosa*’s latent environmental space, that is the effective similarity among distinct environments from the perspective of *P. aeruginosa* genome evolution and therefore indicates the potential ease of ecological transitions across environments through pre-adaptation or shared environmental filtering.

### Environmental source drives phenotypic convergence

Our comparative genomic analyses support the CCS model by revealing both a shallow phylogenetic association with environment (Fig. 1) and consistent genomic associations with environmental source across *P. aeruginosa* strains (Figs. 2-3). These associations are consistent with both environmental filtering (genomic features required for establishment in an environment) and ongoing convergent adaptation within an environment.

Testing our predictions using publicly available genomic databases provides powerful scale (6,627 high-quality genomes with matching environmental metadata, Figs. 1-3) but does not directly address a core implication of the CCS model: strain phenotypes will be shaped more by their environment than by their common ancestry. To test this phenotypic prediction, we assayed an environmentally diverse library of 47 sequenced clade A strains sourced from acute infections, the lungs of patients with cystic fibrosis, and environmental non-human sources for three complementary phenotypic traits relevant to CCS: growth rate, as an integrative proxy for metabolic adaptation to nutrient conditions^11,38^; biofilm formation, a key surface attachment and persistence trait linked to top-ranked CCS genes like *mucA*^39,40^; and protease production, a virulence-associated and environmentally responsive trait, under control of top ranked CCS gene *lasR*^41,42^) (Fig. 4a,b). We then used partial redundancy analysis (partial RDA) to test whether environment explains multivariate phenotypic variation after accounting for phylogenetic structure. Environment explained a significant unique fraction of phenotypic variation after controlling for phylogeny (F = 3.81, p = 0.009), whereas the unique contribution of phylogeny was marginal (F = 1.45, p = 0.091), consistent with the PCA structure (Fig. 4b). PERMDISP did not detect significant differences in within-group dispersion for environment (p = 0.73), suggesting that the environment-associated signal is unlikely to be driven primarily by unequal dispersion.

**Figure 4.**
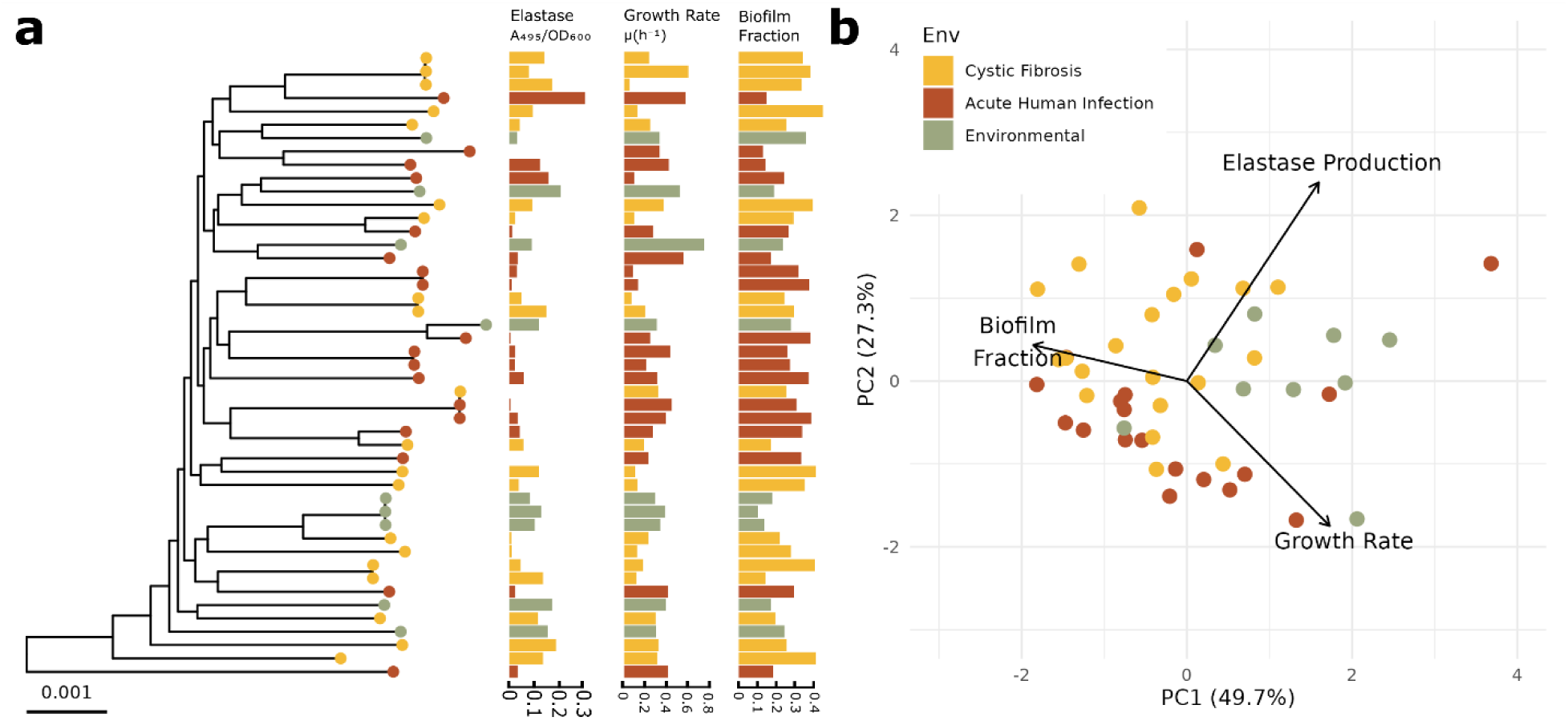
Experimental phenotypes cluster more strongly by environment than by phylogeny. **(a)** Maximum-likelihood phylogeny of the 47-strain experimental library with three phenotypic measurements, growth rate, biofilm formation, and elastase production, plotted alongside each tip (see methods for phenotype measurement details). Phenotypic values show limited phylogenetic structure, with closely related strains often exhibiting divergent trait profiles. **(b)** Principal component analysis (PCA) of phenotypic data. Samples are colored by environment, with loading vectors showing relative contributions of individual phenotypic traits to PC1 and PC2 (longer arrows = stronger influence). The first two principal components captured 77% of the phenotypic variation. Partial RDA confirms that environment explains a significant unique fraction of phenotypic variation after controlling for phylogeny (*F =* 3.81*, p =* 0.009), whereas phylogeny contributes only marginally (*F =* 1.45*, p =* 0.091).

## Discussion

This study challenges the common portrayal of *Pseudomonas aeruginosa* as a strict generalist, and instead supports convergent, cryptic specialization (CCS) due to reproducible, phylogenetically-independent genotype–environment associations. Across >6,600 genomes, environments of isolation are broadly dispersed across the phylogeny, inconsistent with strict lineage-locked specialism, yet genome content predicts environment well above chance under lineage-blocked evaluation. Interpretable feature profiles (SHAP) and complementary mutation-spectrum analyses indicate consistent, non-random shifts in key loci, particularly across chronic lung environments, while phenotyping of a diverse strain panel shows that CCS-relevant phenotypes align more strongly with environmental source than with phylogenetic relatedness. Together, these results suggest that apparent generalism can mask mixtures of cryptic specialists shaped by shared ecological filters and/or convergent selection.

A striking pattern emerges within chronic lung infections. Adult CF, pediatric CF, and non-CF bronchiectasis share many predictive features, implying broadly similar selective pressures across chronic lung disease, yet remain distinguishable by a smaller set of environment-specific variants. The consistent enrichment of non–wild-type *mucA* across chronic lung environments supports a general role for mucoidy in chronic persistence. At the same time, pediatric CF retains distinct markers (e.g., *tonB2*), and bronchiectasis is distinguished by signatures including putative loss-of-function in *phzD1* and other phenazine-associated genes, suggesting additional environment- or stage-specific pressures layered onto a shared chronic-lung background. These differences motivate the view that chronic lung infections share a common adaptive “core” while retaining disease-context specificity.

Several limitations temper inference about mechanism. First, our ecological labels and sampling are constrained by public metadata and clinical collection practices, which may bias the observed frequency of genotypes and blur boundaries among categories. Second, above-chance predictability demonstrates reproducible genomic structure but does not by itself establish causality for any particular locus. The observed genotype–environment associations could arise because certain variants increase establishment success (environmental filtering), and/or because variants are favored after colonization (within-environment evolution), and their balance likely varies across environments. Third, some environments may be under-sampled or too coarsely defined to yield stable contrasts, and misclassifications likely reflect ecological overlap, label noise, and/or smaller class sizes in acute infection categories.

CCS offers a powerful framework for interpreting how a single bacterial species can occupy diverse non-human reservoirs and clinical contexts without deep lineage partitioning, offering a new solution to the paradox of generalism. Our work repositions *P. aeruginosa* as a genomically flexible multi-environment opportunist rather than a strict specialist or generalist. More broadly, our data-led “genotype-discernible environment” approach provides a scalable way to map ecological structure from genomes, prioritize candidate loci and pathways for mechanistic follow-up, and identify representative strains for experimental study (for example, identifying distinct pediatric and adult CF experimental model strains). Perhaps most importantly, our results demonstrate that strains of so-called “generalist” bacterial species can differ significantly and repeatedly across multiple environments, with important implications for both ecology and human health; some strains from particular environments may pose greater systematic risks to humans with defined vulnerabilities (e.g. lung versus wound) than others.

## Methods

### Genome Dataset Assembly

We downloaded metadata for all genomes matching *Pseudomonas aeruginosa* from NCBI as of June 6th 2024 using the NCBI Datasets package^43^, yielding 30,949 candidate samples. Genomes were initially filtered for quality (N50 > 100,000; contig number < 300) and assigned isolation source using Assembly. BioSample.Isolation. source and Assembly. BioSample.Host.disease according to defined matching rules (described in Supplementary Information). Pediatric CF samples were distinguished from adult CF by filtering on patient age (<20 years) where available. Samples annotated only as “sputum” were excluded due to ambiguity across CF and non-CF contexts. Genomes with usable metadata (11,617) were downloaded using NCBI Datasets, and assembly quality was assessed with CheckM^44^; genomes were retained if completeness was >98% and contamination <2%. Pairwise average nucleotide identity (ANI) was computed using fastANI^45^, and genomes were grouped into genomovars (99.5% ANI) and clones (99.99% ANI)^15^. Assembly submitter metadata was retained for downstream bias analyses. All genomes were re-annotated in a single batch using Bakta^46^ to ensure consistency.

### Data Analysis

All statistical analyses and visualization were produced using R version 4.5.2^47^ with ‘tidyvers’^48^. Figures were produced using ‘ggplot2’^49,^ ‘patchwork’^50^, and, for phylogenetic trees, ‘ggtre’^51^ and ‘ggtreeExtrà^52^. Statistical analysis used ‘vegan’^53^, ‘ap’^54^, ‘lme4’^55^, ‘emmeans’^56^ and the built in ‘stats’ package.

### Phylogenetic Reconstruction

A maximum likelihood phylogeny was constructed for the 11,401 genomes passing the above thresholds using Parsnp^57^, with PA7 (GCA_000017205.1) as the outgroup. The tree recapitulated expected population structure (Supplementary Fig. 1), resolving into two major clades (A and B) and a smaller divergent lineage corresponding to *P. paraeruginosa* (formerly clade C)^58^. Clade assignment was performed by reciprocal ANI comparison to reference strains PAO1 (clade A), PA14 (clade B), and PA7 (clade C), assigning genomes based on highest ANI. Clade C genomes (n = 38) were removed, except for PA7 retained for rooting. Due to deep divergence between clades A and B (Supplementary Fig. 1), lower sample size and lower environmental heterogeneity in clade B, as well as an excess of closely related isolates, clade B genomes (n = 2,771) were excluded to avoid confounding model performance. All downstream analyses were restricted to clade A, though model performance (trained on clade A) was evaluated on clade B (Supplementary Fig 2).

### Trait Depth Analysis

Phylogenetic depth of environment-of-origin associations was quantified using the consenTRAIT algorithm^14^. For each environment, consenTRAIT identifies the deepest internal node whose descendant tips share the trait in at least 90% of members and computes 𝜏_𝐷_ as the mean branch length from that node to associated tips. Singleton occurrences were assigned a depth equal to half the distance to the nearest internal node to maintain comparability across traits. To evaluate whether observed environmental clustering exceeded random expectation, trait labels were permuted across the phylogeny to generate a null distribution of 𝜏_𝐷_values, and observed values were compared with this null distribution for significance testing. Mean 𝜏_𝐷_values for each environment were mapped onto the phylogeny to visualize the depth and distribution of environment-specific associations.

### Genomovar and ANI-Based Clustering

Strains from Clade A were clustered into increasingly fine-scale phylogenetic groups using pairwise average nucleotide identity (ANI) thresholds from 99.3% to 99.99%. These thresholds capture a hierarchy of relatedness, from broad subclade structure to clonal groups. Genomovars were defined at ANI >99.5%, consistent with established conventions^15^. For each ANI threshold, strains were assigned to non-overlapping clusters, and the diversity of environments of isolation represented within each cluster was quantified to assess how environmental breadth changed with phylogenetic resolution. Clusters containing fewer than nine strains were excluded from diversity calculations to limit bias from sparse sampling.

### Feature Generation

In order to distinguish protein-coding variants across the 6,627 genomes included in the study, we modified the source code of Panaroo^18^ to enable exact protein-variant grouping and classification. Panaroo normally clusters proteins through successive rounds of sequence similarity-based grouping; we altered this behavior to require 100% protein identity and 100% protein-length match and removed lower-threshold regrouping steps. Using these settings, Panaroo identified 1,337,690 unique protein variants across the 11,598 genomes that passed quality filtering, including distinct genes and coding variants differing by as little as a single amino acid. We then used ‘gene_presence_absence.Rtab’ to encode presence/absence of each protein variant and filtered out variants with <0.1% or >99.9% prevalence, as well as variants that showed perfect co-segregation across genomes. After feature generation, we removed samples with duplicate feature patterns, which corresponded to duplicate genomes (161 samples).

### XGBoost Model Implementation

Supervised machine learning was used to test whether genomic features predict environment of origin. Under strict generalism (H2), model performance should be comparable to randomized baselines; under convergent cryptic specialization (H3), performance should exceed baseline. An 80:20 train-validation split was performed before any model fitting, and the validation set was held out until final evaluation. XGBoost^17^ models were trained to predict environment of origin from the presence and absence of exact protein-coding variants. XGBoost was chosen because tree-based models provide interpretable decisions on binary features and are less prone to overfitting in high-dimensional, low-sample settings. Model training was controlled through configuration files specifying the label group, labels to exclude, feature count, and hyperparameter-tuning strategy. For initial blocked cross-validation models, the feature set was limited to 20,000 variables using ANOVA F-scores. Hyperparameters were optimized by Bayesian search over five rounds. In each round, a baseline model was initialized, the training data were split into five stratified folds, models were fit on four folds and evaluated on the held-out fold, and median accuracy across folds was used to score the parameter set. The process was repeated for the specified number of rounds.

### Genomovar Blocked Cross-Validation Strategy

Because phylogenetic structure in environment-of-origin was detectable only weakly below the genomovar level (99.5% ANI), we used genomovar-blocked cross-validation. In each fold, all samples from a single genomovar were withheld while the XGBoost model was trained on all other genomovars; performance was then evaluated on the withheld genomovar, and SHAP^20^ values were computed for those predictions. This approach limits phylogenetic confounding by ensuring that the model never observes the withheld genomovar during training. The procedure was repeated for every genomovar and every environment contrast, yielding out-of-genomovar predictions and genomovar-resolved feature-importance profiles.

### Identification of Genotype-Discernible Environments

XGBoost genomovar-blocked cross-validation models were first trained on all 17 metadata-defined environments. Class-wise performance was quantified using normalized accuracy, defined as observed accuracy minus stratified random-guessing accuracy. Environments with normalized accuracy consistently below zero were considered indistinguishable from chance and were evaluated for possible merging with ecologically similar categories (e.g., Ocean with Aquatic). When a low-performing environment had a plausible merge partner, the merged category was re-evaluated in a new round of genomovar-blocked cross-validation. If the merged category performed worse than the original or if performance remained low after merging, the environment(s) were removed. Regrouped samples kept their original train-test split to preserve the stratified label distribution. The “Animal” environment was removed due to low performance (<10% maximum normalized accuracy) and small sample size, though more data might enable better separation. “Pneumonia” was excluded as an independent category because it degraded overall performance despite low accuracy (∼6%). The “Lung” environment, defined by low-resolution terms such as “infected lung,” “respiratory tract infection,” or “bronchoalveolar lavage,” was removed because the underlying source (e.g., CF, non-CF bronchiectasis, or pneumonia) could not be resolved. The final retained set comprised nine environments (Adult CF, Pediatric CF, non-CF bronchiectasis, blood, urinary, fecal, wound, Aquatic, and hospital) that consistently exceeded the stratified chance baseline across resampling.

### Feature Filtering

To prepare the feature set for final evaluation of the nine-environment model on held-out validation data, we applied sequential filtering steps that enriched for broadly common and informative protein-coding variants while removing those likely to reflect study-specific artifacts or narrow genomovar distributions. We started with 2,773 protein-coding variants that had SHAP values across all blocked-cross-validation folds. First, we removed variants flagged as study-specific, defined as those with >50% relative-abundance difference between the highest- and second-highest-abundance studies. Next, we retained only variants selected in all cross-validation blocks. Finally, we required sufficient predicted-state prevalence across genomovars: predicted-present variants had to appear in ≥25% of genomovars and predicted-absent variants had to be absent in ≥25% of genomovars. This process yielded 477 variants for final evaluation.

### Model Evaluation

Trained models were evaluated on the test split (held out validation dataset) using the final 9 environment groups derived during training. Performance was assessed using micro-F1, macro-F1, accuracy, and balanced accuracy. By default, models were trained with a multiclass softmax objective; alternatively, a multiclass softprob objective was used to obtain one-vs-all ROC metrics.

### SHAP Scoring and Functional Annotation

SHAP (SHapley Additive exPlanations) is a game-theoretic approach to model interpretability that quantifies the contribution of each feature to an individual prediction^20^. SHAP treats the model output as the sum of feature effects, evaluating how each feature value shifts the prediction relative to a baseline. For genomic data, SHAP computes the effect of a gene’s presence or absence on the predicted class logit or probability. Each feature is assigned a SHAP value representing the magnitude and direction of its contribution to a given prediction; large absolute values indicate strong influence. The sign indicates whether the feature pushes the model toward (positive) or away from (negative) a particular class. SHAP values were used to interpret how the environment-predicting XGBoost models made their predictions across the nine retained environments.

### Variant-Level Predictive Contribution

We used two complementary SHAP-based summaries: a ΔSHAP statistic for gene-level, environment-specific interpretation, and an environment-similarity metric in ENV space.

For each gene 𝑔 and environment 𝑒, SHAP values 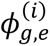 were computed for all isolates 𝑖, and the contrast

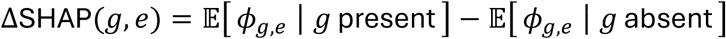

was used to quantify whether the presence of 𝑔 tends to increase or decrease the model’s support for 𝑒 relative to its absence. This retains both magnitude and direction, whereas standard mean absolute SHAP discards sign and can swamp environment-specific effects. For environment-environment similarity, we avoided pure mean absolute SHAP and instead defined each gene’s expected state (present or absent) per environment, then summarized that gene’s contribution to each environment by the mean SHAP under the expected state. Environment–environment similarity was computed by aggregating absolute values of these expected-state contributions across genes, so that environments are considered similar when they are supported by the same genes in the same expected states with comparable strength. Both ΔSHAP and the ENV-similarity metric are explicitly directional and environment-specific, aligning the summaries with the underlying biological questions rather than generic feature importance.

### Environment-Level Similarity and Network Analysis

A network was constructed in which nodes represent environments and edges represent the total shared SHAP-based contribution of protein-coding variants with concordant directionality between pairs of environments. Edge weights were computed from pairwise sums of SHAP or net-predictive-value contributions across shared genes, restricted to non-zero edges, and the graph was implemented using igraph^59^ to visualize and assess similarity among chronic lung, acute infection, and non-host environments.

### Variant Mutation Analysis

We used reference genes from *P. aeruginosa* PAO1 (NC_002516.2) to categorize CF-adaptive variants as wild-type or mutant. Genes were identified in all genomes using nucleotide BLAST; missing genes were labelled as deleted and those split across multiple contigs or carrying insertions as incomplete. Gene regions were extracted with bedtools2 getfasta and translated with EMBOSS transeq; protein sequences were then compared to PAO1 using protein BLAST to compute query coverage (alignment length × percent identity / reference length). Proteins with 100% coverage were classified as wild-type (functional), those with ≤40% coverage as non-functional, and those in between were clustered by CD-Hit^60^ at 100% identity and length, then aligned to references with Clustal Omega^61^. Amino acid substitutions, frameshifts, premature stop codons, insertions, and deletions were extracted from the alignments using Biopython^62^; proteins with frameshifts or premature stops, plus previously identified deleted and incomplete genes, were assigned “no function,” whereas those with amino acid substitutions were labelled “alternate function.” Genes were classified as “contig splits” if BLAST^63^ yielded a partial match on a different contig that included the start or stop of the contig (within 5 bp, applying this window only to the stop to account for mismatches), or if the gene had multiple partial matches across more than one contig.

### Mutation Status Statistical Analysis

For genes of interest, we fit binomial generalized linear mixed models (GLMMs) separately for each gene with mutation status (eg. functional vs. alternative function) as the binary response, environment of isolation as a fixed effect, and genomovar group as a random intercept to control for phylogenetic structure.

Estimated marginal means of mutation probabilities were computed for each niche using emmeans^56^ with Tukey-adjusted pairwise contrasts^47^. Plots display marginal probabilities with 95% confidence intervals and compact letter display notation indicating significant differences (P < 0.05). Analyses were repeated (1) excluding contig-split mutations and (2) restricting to loss-of-function mutations only.

### Phenotypic Strain Library

An environmentally diverse library of 47 sequenced strains from Clade A was used to measure growth rate, biofilm formation, and protease (elastase) production.

### Media

Unless otherwise stated, experiments were performed in a defined rich medium (hereafter “growth medium”) consisting of M9 salts supplemented with 10 mM glucose, 1% casamino acids, 1 mM MgSO₄, 100 µM CaCl₂, and 1× Hutner’s trace elements^64^.

### Growth curves

Mid-exponential phase cultures were diluted to a starting optical density (OD₆₀₀) of 0.05 in 150 µL of growth medium in 96-well microtiter plates. Plates were incubated at 37 °C with continuous orbital shaking in a Varioskan plate reader, with OD₆₀₀ measured hourly to capture growth dynamics. Growth curves were modeled using GrowthCruvR^65^, and the maximum growth rate (𝜇) was extracted for each strain from the fitted logistic curves.

### Biofilm formation and composition

Cultures were diluted to OD₆₀₀ = 0.05 in 150 µL growth medium and incubated statically in 96-well plates at 37 °C for 6 h. Planktonic OD₆₀₀ was measured, the supernatant was removed, and biofilms were washed with sterile water. Biofilms were resuspended in 150 µL 0.85% NaCl using 1 mm glass beads and vortexing (15 min, 2000rpm), then transferred to a new plate for a resuspended biofilm OD₆₀₀ measurement. To quantify the proportion of total biomass allocated to the biofilm, the biofilm fraction was calculated by dividing the resuspended biofilm OD₆₀₀ by the sum of the resuspended biofilm OD₆₀₀ and the planktonic OD₆₀₀.

### Elastase assay

Cultures were grown in 150 µL growth medium at 37 °C with shaking and harvested at late exponential phase based on expected transitions from prior growth curves. At the appropriate time, OD₆₀₀ was recorded, cultures were transferred to 1.5 mL tubes, and cells were pelleted (14,000 rpm, 1 min). Supernatants were filter-sterilized (0.22 µm) to obtain cell-free supernatant, and downstream analyses were performed immediately.

Elastase activity was measured using the elastin–Congo red (ECR) assay (Ohman et al., 1980). Briefly, 100 µL of cell-free supernatant was added to 900 µL ECR buffer (100 mM Tris, 1 mM CaCl₂, pH 7.5) containing 20 mg ECR (Sigma) and incubated at 37 °C with shaking for 20 h. Remaining insoluble substrate was pelleted via centrifugation (14,000 rpm, 1 min), and 200 µL of the resulting supernatant was transferred to a 96-well plate. Absorbance was measured at 495 nm, and values were reported as normalized to the OD₆₀₀ of the culture at the time of harvest.

### Phylogenetic reconstruction and variance partitioning for phenotypic experiments

A maximum likelihood phylogeny was constructed for the 47 strains used in phenotypic assays using Parsnp^57^, with PA7 (GCA_000017205.1) as the outgroup. The tree was rooted on PA7, which was subsequently removed prior to downstream analyses.

Phylogenetic structure was represented by eigenvectors derived from the cophenetic distance matrix. Distances were projected into Euclidean space using classical multidimensional scaling with additive correction, retaining only axes with positive eigenvalues. The minimal set of axes explaining 80% of this variance (maximum 10 axes) was used as phylogenetic predictors. Environment was encoded as a model matrix, and variance partitioning was performed using partial redundancy analysis (RDA), with significance assessed by 999 permutations.

## Reference

1. Poisot, T., Bever, J. D., Nemri, A., Thrall, P. H. & Hochberg, M. E. A conceptual framework for the evolution of ecological specialisation. Ecol Lett 14, 841–851 (2011).

2. Schick, A., Bailey, S. F. & Kassen, R. Evolution of Fitness Trade-Offs in Locally Adapted Populations of Pseudomonas fluorescens. Am Nat 186 Suppl 1, S48–59 (2015).

3. MacArthur, R. H. & Pianka, E. R. On Optimal Use of a Patchy Environment. The American Naturalist 100, 603–609 (1966).

4. Levins, R. Theory of Fitness in a Heterogeneous Environment. II. Developmental Flexibility and Niche Selection. The American Naturalist 97, 75–90 (1963).

5. Bono, L. M., Smith, L. B., Pfennig, D. W. & Burch, C. L. The emergence of performance trade-offs during local adaptation: insights from experimental evolution. Mol Ecol 26, 1720–1733 (2017).

6. Kassen, R. The experimental evolution of specialists, generalists, and the maintenance of diversity. Journal of Evolutionary Biology 15, 173–190 (2002).

7. Buckling, A., Wills, M. A. & Colegrave, N. Adaptation Limits Diversification of Experimental Bacterial Populations. Science 302, 2107–2109 (2003).

8. Sriswasdi, S., Yang, C. & Iwasaki, W. Generalist species drive microbial dispersion and evolution. Nat Commun 8, 1162 (2017).

9. Winstanley, C., O’Brien, S. & Brockhurst, M. A. Pseudomonas aeruginosa Evolutionary Adaptation and Diversification in Cystic Fibrosis Chronic Lung Infections. Trends Microbiol 24, 327–337 (2016).

10. Lor’, N. I. et al. Cystic fibrosis-niche adaptation of Pseudomonas aeruginosa reduces virulence in multiple infection hosts. PLoS One 7, e35648 (2012).

11. La Rosa, R., Johansen, H. K. & Molin, S. Convergent Metabolic Specialization through Distinct Evolutionary Paths in Pseudomonas aeruginosa. mBio 9, e00269–18 (2018).

12. Marvig, R. L., Sommer, L. M., Molin, S. & Johansen, H. K. Convergent evolution and adaptation of Pseudomonas aeruginosa within patients with cystic fibrosis. Nat Genet 47, 57–64 (2015).

13. Smith, E. E. et al. Genetic adaptation by Pseudomonas aeruginosa to the airways of cystic fibrosis patients. Proceedings of the National Academy of Sciences 103, 8487–8492 (2006).

14. Martiny, A. C., Treseder, K. & Pusch, G. Phylogenetic conservatism of functional traits in microorganisms. ISME J 7, 830–838 (2013).

15. Rodriguez-R, L. M. et al. An ANI gap within bacterial species that advances the definitions of intra-species units. mBio 15, e02696–23 (2023).

16. Jain, C., Rodriguez-R, L. M., Phillippy, A. M., Konstantinidis, K. T. & Aluru, S. High throughput ANI analysis of 90 K prokaryotic genomes reveals clear species boundaries. Nat Commun 9, 5114 (2018).

17. Chen, T. & Guestrin, C. XGBoost: A Scalable Tree Boosting System. in Proceedings of the 22nd ACM SIGKDD International Conference on Knowledge Discovery and Data Mining 785–794 (Association for Computing Machinery, New York, NY, USA, 2016). doi:10.1145/2939672.2939785.

18. Tonkin-Hill, G. et al. Producing polished prokaryotic pangenomes with the Panaroo pipeline. Genome Biol 21, 180 (2020).

19. Roberts, D. R. et al. Cross-validation strategies for data with temporal, spatial, hierarchical, or phylogenetic structure. Ecography 40, 913–929 (2017).

20. Lundberg, S. M. & Lee, S.-I. A Unified Approach to Interpreting Model Predictions. in Advances in Neural Information Processing Systems vol. 30 (Curran Associates, Inc., 2017).

21. Martin, D. W. et al. Mechanism of conversion to mucoidy in Pseudomonas aeruginosa infecting cystic fibrosis patients. Proc Natl Acad Sci U S A 90, 8377–8381 (1993).

22. Wood, L. F. & Ohman, D. E. Identification of Genes in the σ22 Regulon of Pseudomonas aeruginosa Required for Cell Envelope Homeostasis in Either the Planktonic or the Sessile Mode of Growth. mBio 3, 10.1128/mbio.00094-12 (2012).

23. Bhagirath, A. Y. et al. Cystic fibrosis lung environment and Pseudomonas aeruginosa infection. BMC Pulm Med 16, 174 (2016).

24. Schalk, I. J. & Perraud, Q. Pseudomonas aeruginosa and its multiple strategies to access iron. Environmental Microbiology 25, 811–831 (2023).

25. Tralau, T. et al. Transcriptomic Analysis of the Sulfate Starvation Response of Pseudomonas aeruginosa. Journal of Bacteriology 189, 6743–6750 (2007).

26. Llanes, C. et al. Clinical Strains of Pseudomonas aeruginosa Overproducing MexAB-OprM and MexXY Efflux Pumps Simultaneously. Antimicrob Agents Chemother 48, 1797–1802 (2004).

27. Laborda, P. et al. Mutations in the efflux pump regulator MexZ shift tissue colonization by Pseudomonas aeruginosa to a state of antibiotic tolerance. Nat Commun 15, 2584 (2024).

28. Liang, X. et al. Identification of divergent type VI secretion effectors using a conserved chaperone domain. Proc Natl Acad Sci U S A 112, 9106–9111 (2015).

29. Pan, X. et al. Pseudomonas aeruginosa lasR-deficient mutant contributes to bacterial virulence through enhancing the PhoB-mediated pathway in response to host environment. mBio 16, e01788–25 (2025).

30. Zhao, K. et al. Evolution of lasR mutants in polymorphic Pseudomonas aeruginosa populations facilitates chronic infection of the lung. Nat Commun 14, 5976 (2023).

31. Gambello, M. J., Kaye, S. & Iglewski, B. H. LasR of Pseudomonas aeruginosa is a transcriptional activator of the alkaline protease gene (apr) and an enhancer of exotoxin A expression. Infection and Immunity 61, 1180–1184 (1993).

32. Irby, I., Mehlferber, E. C. & Brown, S. P. Canonical cystic fibrosis genes in P. aeruginosa are not CF specific and are enriched across chronic lung infections. bioRxiv.

33. Mavrodi, D. V. et al. Functional analysis of genes for biosynthesis of pyocyanin and phenazine-1-carboxamide from Pseudomonas aeruginosa PAO1. J Bacteriol 183, 6454–6465 (2001).

34. Schiessl, K. T. et al. Phenazine production promotes antibiotic tolerance and metabolic heterogeneity in Pseudomonas aeruginosa biofilms. Nat Commun 10, 762 (2019).

35. Mavrodi, D. V. et al. Functional Analysis of Genes for Biosynthesis of Pyocyanin and Phenazine-1-Carboxamide from Pseudomonas aeruginosa PAO1. Journal of Bacteriology 183, 6454–6465 (2001).

36. O’Connor, K., Zhao, C. Y., Mei, M. & Diggle, S. P. Frequency of quorum-sensing mutations in Pseudomonas aeruginosa strains isolated from different environments. Microbiology 168, 001265 (2022).

37. Trottier, M. C. et al. The end of the reign of a “master regulator’’? A defect in function of the LasR quorum sensing regulator is a common feature of Pseudomonas aeruginosa isolates. mBio 15, e02376–23 (2024).

38. Frimmersdorf, E., Horatzek, S., Pelnikevich, A., Wiehlmann, L. & Schomburg, D. How Pseudomonas aeruginosa adapts to various environments: a metabolomic approach. Environmental Microbiology 12, 1734–1747 (2010).

39. Xie, Z. D., Hershberger, C. D., Shankar, S., Ye, R. W. & Chakrabarty, A. M. Sigma factor-anti-sigma factor interaction in alginate synthesis: inhibition of AlgT by MucA. Journal of Bacteriology 178, 4990–4996 (1996).

40. Wang, Y.-H. et al. Mutations in the mucA gene of Pseudomonas aeruginosa promote disease exacerbation. Arch Microbiol 208, 200 (2026).

41. Le Berre, R. et al. Quorum-sensing activity and related virulence factor expression in clinically pathogenic isolates of *Pseudomonas aeruginosa*. Clinical Microbiology and Infection 14, 337–343 (2008).

42. Toder, D. S., Gambello, M. J. & Iglewski, B. H. Pseudomonas aeruginosa LasA: a second elastase under the transcriptional control of lasR. Molecular Microbiology 5, 2003–2010 (1991).

43. O’Leary, N. A. et al. Exploring and retrieving sequence and metadata for species across the tree of life with NCBI Datasets. Sci Data 11, 732 (2024).

44. Parks, D. H., Imelfort, M., Skennerton, C. T., Hugenholtz, P. & Tyson, G. W. CheckM: assessing the quality of microbial genomes recovered from isolates, single cells, and metagenomes. Genome Research 25, 1043–1055 (2015).

45. Jain, C., Rodriguez-R, L. M., Phillippy, A. M., Konstantinidis, K. T. & Aluru, S. High throughput ANI analysis of 90 K prokaryotic genomes reveals clear species boundaries. Nat Commun 9, 5114 (2018).

46. Schwengers, O. et al. Bakta: rapid and standardized annotation of bacterial genomes via alignment-free sequence identification. Microb Genom 7, 000685 (2021).

47. R Core Team. R: A Language and Environment for Statistical Computing. (R Foundation for Statistical Computing, Vienna, Austria, 2026).

48. Wickham, H. et al. Welcome to the tidyverse. Journal of Open Source Software 4, 1686 (2019).

49. Wickham, H. Ggplot2: Elegant Graphics for Data Analysis. (Springer-Verlag New York, 2016).

50. Pedersen, T. L. Patchwork: The Composer of Plots. (2025).

51. Yu, G. Using ggtree to Visualize Data on Tree-Like Structures. Curr. Protoc. Bioinformatics 69, (2020).

52. Xu, S. et al. ggtreeExtra: Compact visualization of richly annotated phylogenetic data. Molecular Biology and Evolution 38, 4039–4042 (2021).

53. Dixon, P. VEGAN, a package of R functions for community ecology. Journal of Vegetation Science 14, 927–930 (2003).

54. Paradis, E. & Schliep, K. ape 5.0: an environment for modern phylogenetics and evolutionary analyses in R. Bioinformatics 35, 526–528 (2019).

55. Bates, D., Mächler, M., Bolker, B. & Walker, S. Fitting Linear Mixed-Effects Models Using lme4. Journal of Statistical Software 67, 1–48 (2015).

56. Lenth, R. V. & Piaskowski, J. Emmeans: Estimated Marginal Means, Aka Least-Squares Means. (2026).

57. Kille, B. et al. Parsnp 2.0: scalable core-genome alignment for massive microbial datasets. Bioinformatics 40, btae311 (2024).

58. Rudra, B., Duncan, L., Shah, A. J., Shah, H. N. & Gupta, R. S. Phylogenomic and comparative genomic studies robustly demarcate two distinct clades of Pseudomonas aeruginosa strains: proposal to transfer the strains from an outlier clade to a novel species Pseudomonas paraeruginosa sp. nov. International Journal of Systematic and Evolutionary Microbiology 72, 005542 (2022).

59. Csárdi, G. et al. Igraph: Network Analysis and Visualization in R. (2026). doi:10.5281/zenodo.7682609.

60. Fu, L., Niu, B., Zhu, Z., Wu, S. & Li, W. CD-HIT: accelerated for clustering the next-generation sequencing data. Bioinformatics 28, 3150–3152 (2012).

61. Sievers, F. et al. Fast, scalable generation of high-quality protein multiple sequence alignments using Clustal Omega. Mol Syst Biol 7, 539 (2011).

62. Cock, P. J. A. et al. Biopython: freely available Python tools for computational molecular biology and bioinformatics. Bioinformatics 25, 1422–1423 (2009).

63. Altschul, S. F., Gish, W., Miller, W., Myers, E. W. & Lipman, D. J. Basic local alignment search tool. J Mol Biol 215, 403–410 (1990).

64. Hutner, S. H., Provasoli, L., Schatz, A. & Haskins, C. P. Some approaches to the study of the role of metals in the metabolism of microorganisms. Proceedings of the American Philosophical Society 94, 152–170 (1950).

65. Sprouffske, K. & Wagner, A. Growthcurver: an R package for obtaining interpretable metrics from microbial growth curves. BMC Bioinformatics 17, 172 (2016).

