## Supplemental Figures for "Apparent generalism in *Pseudomonas aeruginosa* is underpinned by Convergent, Cryptic Specialization"

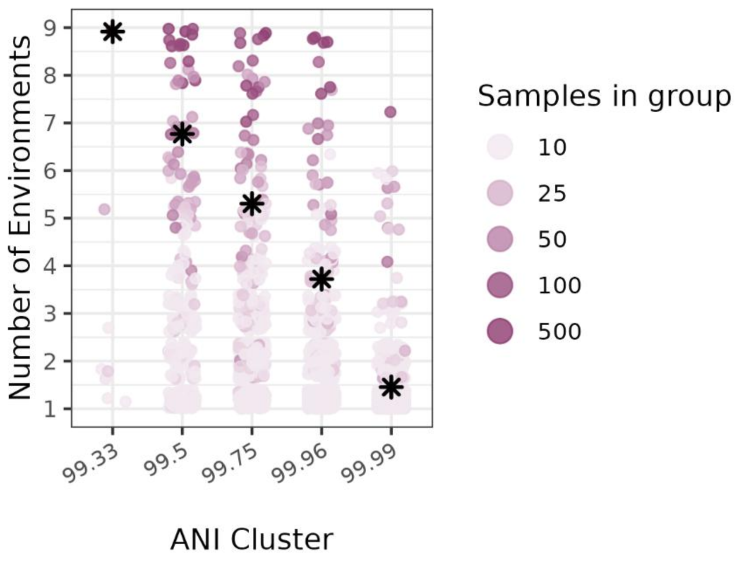


**Supplementary Fig. 1 | Unfiltered ANI clusters.** Environment-of-isolation diversity across hierarchical ANI clusters spanning 99.33%–99.99%, including all clusters regardless of sample size. Results are qualitatively similar to Fig. 1c. Even at the highest ANI thresholds, clusters typically include strains from multiple environments, indicating limited ecological specificity at fine phylogenetic scales. Note that genomovars with less than 9 samples cannot be present in all 9 environments, due to sampling limits alone.

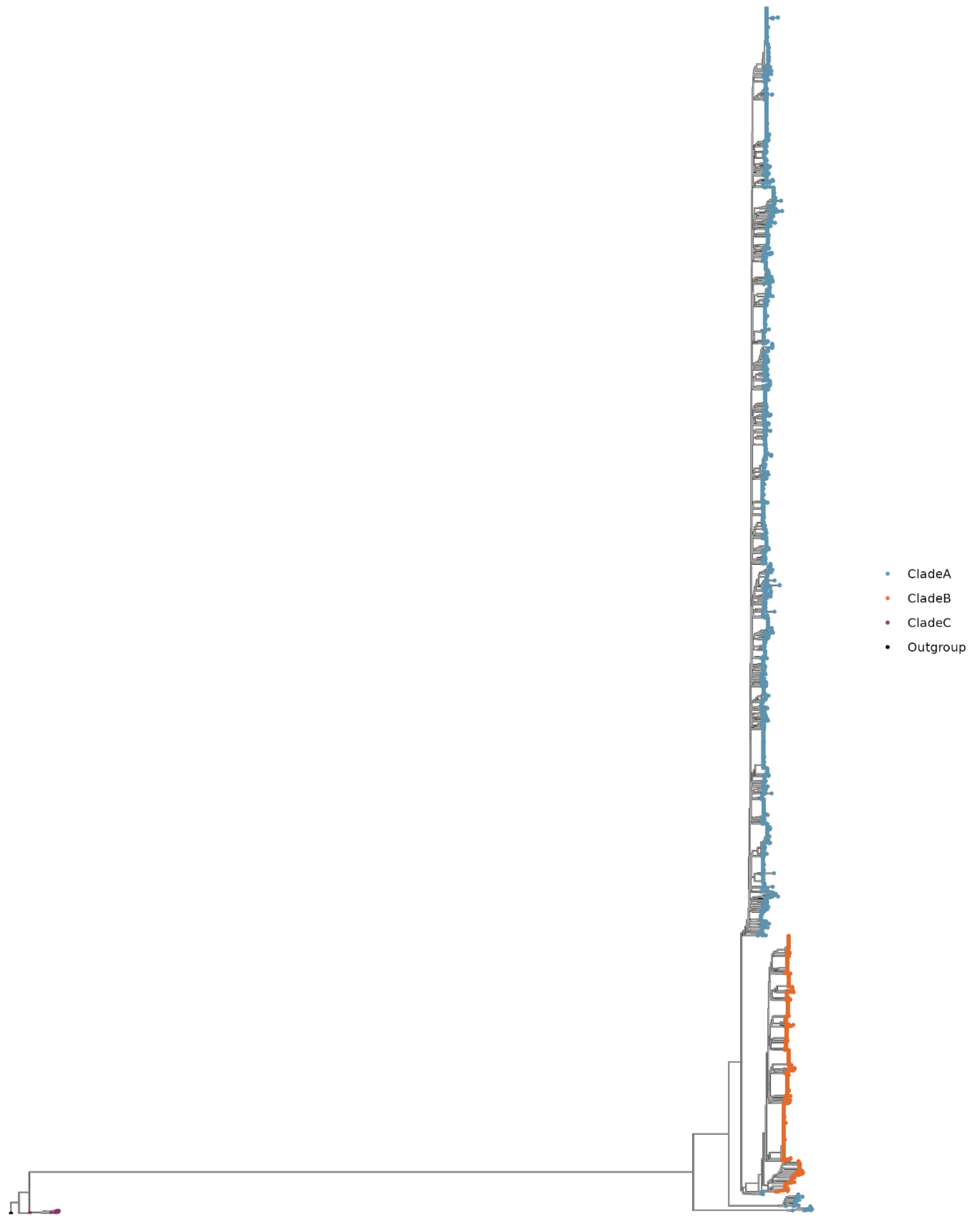


**Supplementary Fig. 2 | Phylogenetic tree of *P. aeruginosa*.** Maximum-likelihood phylogeny of *P. aeruginosa* constructed from 11,401 genomes downloaded from NCBI and rooted with PA7 as the outgroup. Tips are colored by clade assignment (Clade A, blue; Clade B, orange; Clade C, purple), based on reciprocal ANI comparisons to reference strains PAO1 (Clade A), PA14 (Clade B), and PA7 (Clade C), with genomes assigned to the clade showing the highest ANI. Although Clade C has been reclassified as a separate species, *P. paraeruginosa* in NCBI, some strains remain annotated as *P. aeruginosa*; these were excluded from the study. For this paper, subgroups 2 and 3, which fall between *P. aeruginosa* and *P. paraeruginosa*, were treated as part of Clade A on the basis of ANI similarity to PAO1.

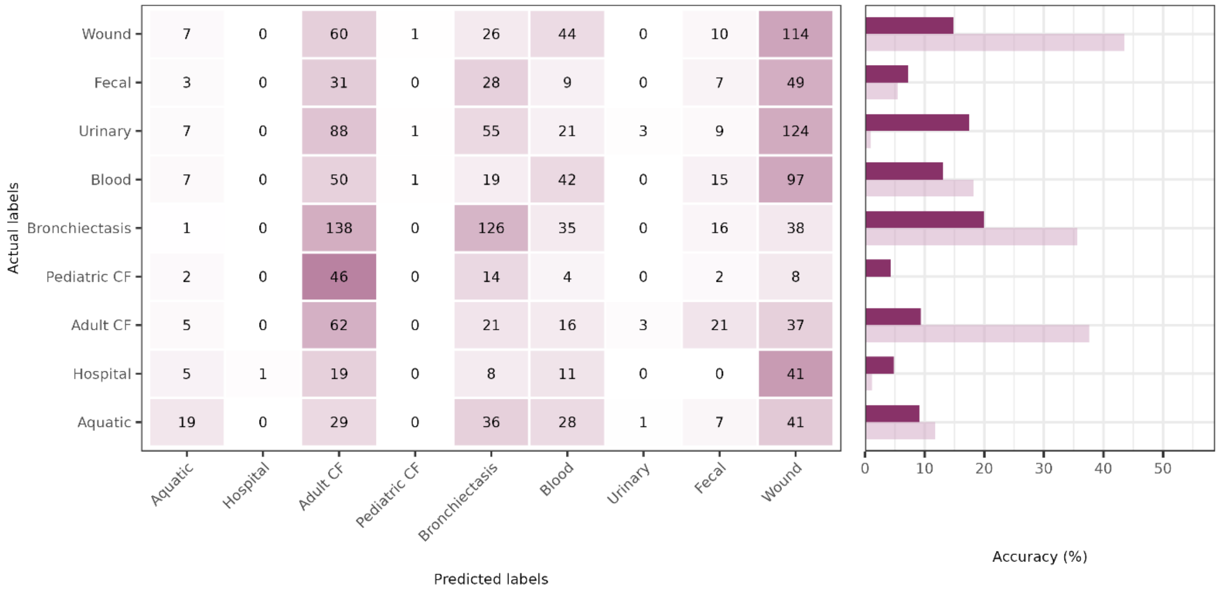


| | Aquatic | Hospital | Adult CF | Pediatric CF | Bronchiectasis | Blood | Urinary | Fecal | Wound |
|---|---|---|---|---|---|---|---|---|---|
| Wound | 7 | 0 | 60 | 1 | 26 | 44 | 0 | 10 | 114 |
| Fecal | 3 | 0 | 31 | 0 | 28 | 9 | 0 | 7 | 49 |
| Urinary | 7 | 0 | 88 | 1 | 55 | 21 | 3 | 9 | 124 |
| Blood | 7 | 0 | 50 | 1 | 19 | 42 | 0 | 15 | 97 |
| Bronchiectasis | 1 | 0 | 138 | 0 | 126 | 35 | 0 | 16 | 38 |
| Pediatric CF | 2 | 0 | 46 | 0 | 14 | 4 | 0 | 2 | 8 |
| Adult CF | 5 | 0 | 62 | 0 | 21 | 16 | 3 | 21 | 37 |
| Hospital | 5 | 1 | 19 | 0 | 8 | 11 | 0 | 0 | 41 |
| Aquatic | 19 | 0 | 29 | 0 | 36 | 28 | 1 | 7 | 41 |

**Supplementary Fig. 3 | Model performance on Clade B.** Performance of the final model trained on 477 filtered features from Clade A. Accuracy is shown at right, with raw accuracy in light pink and random-assignment null accuracy in dark purple. Overall performance was poorer than for Clade A, with several environments performing below random expectation, including fecal, urinary, pediatric CF, and hospital samples (classes the model very rarely predicted). The model performed relatively well on adult CF, bronchiectasis, wound, and blood samples, although many samples were misassigned to these classes. These results suggest that Clade B strains may exhibit a distinct adaptive pattern, particularly in the context of acute infection.

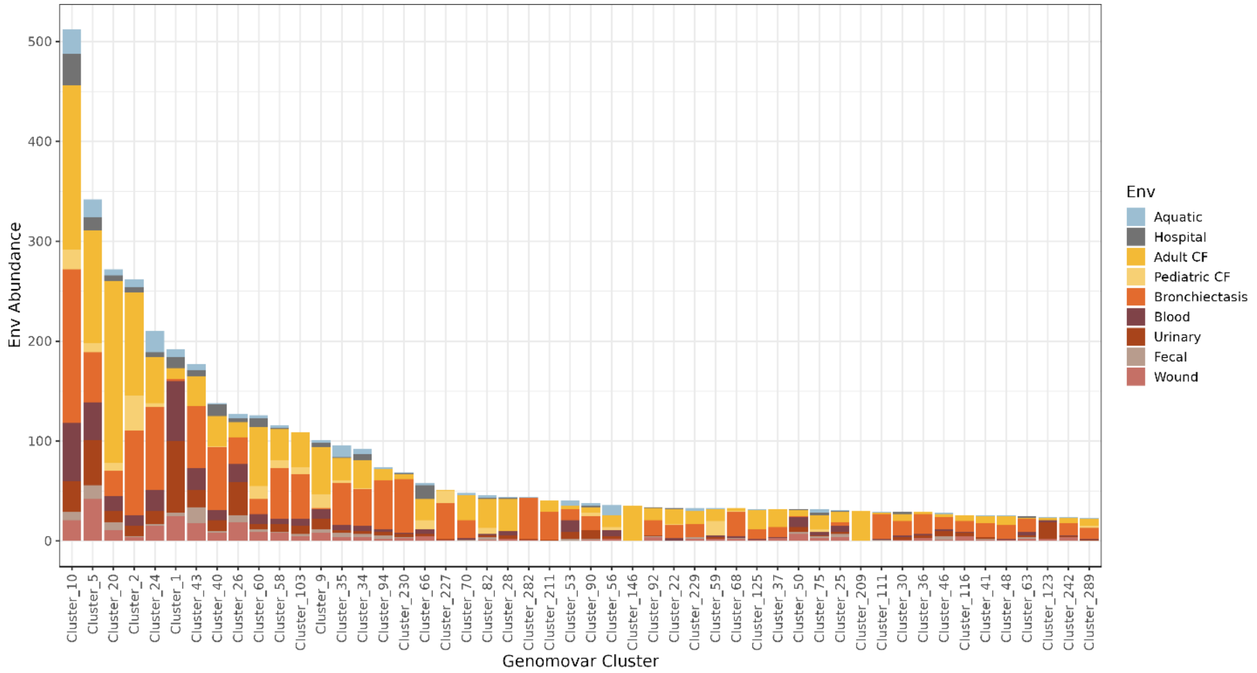


**Supplementary Fig. 4 | Distribution of environments across the top 50 genomovars.** The top 50 Clade A genomovars are shown with their environmental abundances, illustrating the broad distribution of environments among closely related strains.

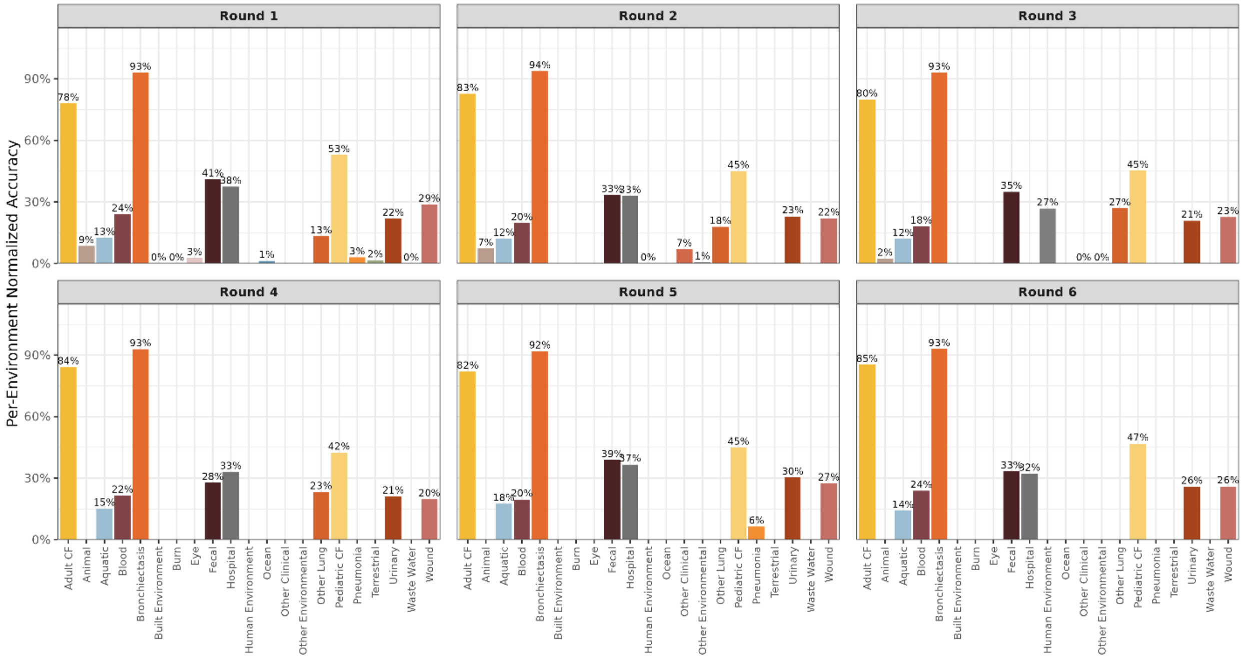


**Supplementary Fig. 5 | Performance of blocked models across candidate environments.** Model performance is shown as normalized accuracy (observed accuracy minus stratified random-guess accuracy) for each environment of origin. The panels summarize the iterative blocked cross-validation approach used to define supported niches for the final model. Environments with normalized accuracy consistently below zero were considered indistinguishable from chance and were evaluated for merging with ecologically similar categories, such as Ocean and Aquatic. When a low-performing environment had a plausible merge partner, the merged category was re-evaluated in a new round of genomovar-blocked cross-validation. If the merged category performed worse than the original or if performance remained low after merging, the environment(s) were removed. The final 9 environment classification is shown in the final (round 6) panel.

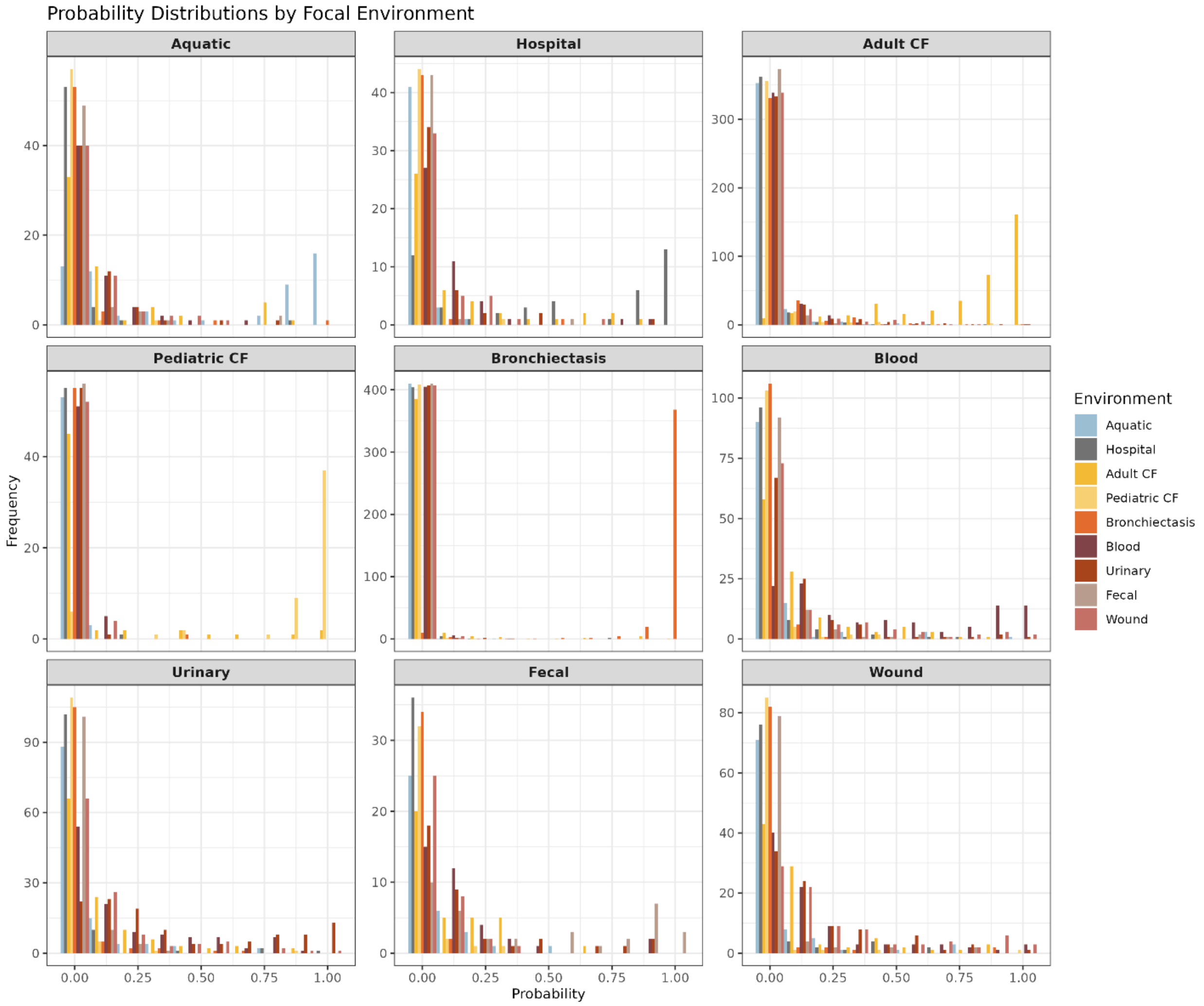


**Supplementary Fig. 6 | Distribution of model environment-assignment probabilities by sample.** The model was trained with a multiclass softprob objective, allowing one-vs-all ROC analysis and visualization of prediction confidence. Each panel shows the frequency and confidence of each assignment for each environment, colored by the true environment of origin for each sample. For the chronic lung environments (Adult CF, Pediatric CF, and Bronchiectasis), the model generally assigns true samples high confidence and rarely assigns high confidence to samples from other environments, indicating confident classification of these groups. In contrast, the acute environments Blood, Urinary, and Wound show more frequent high-confidence misassignments, suggesting overlap in underlying features defining these environments or recent environmental transitions.

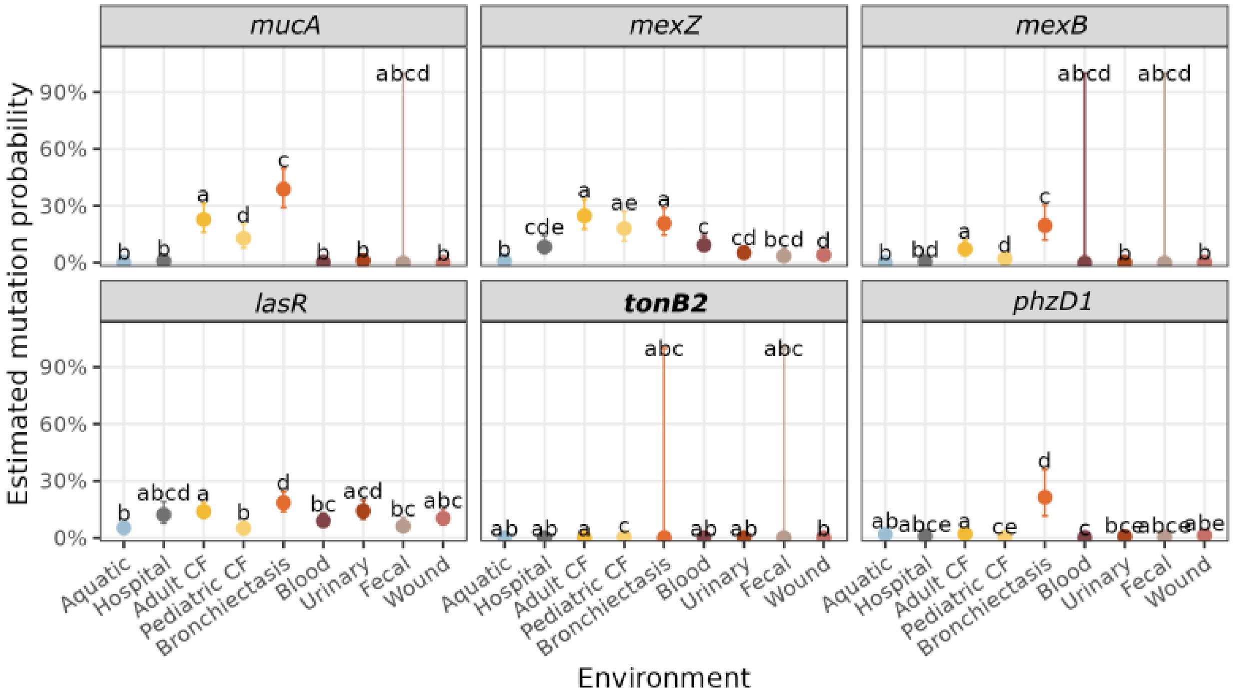


**Supplementary Fig. 7 | Estimated probability of loss-of-function mutation per environment and gene.** Estimated mutation probabilities restricted to putative loss-of-function mutations, from an LMER mixed-effects model, shown as points (± SE) by environment across all *P. aeruginosa* genomes relative to the PAO1 reference (see main text Fig. 3b for same analysis applied to all mutation types). The model includes environment as a fixed effect and genomovar as a random effect, with Tukey post hoc tests used to assess pairwise differences.

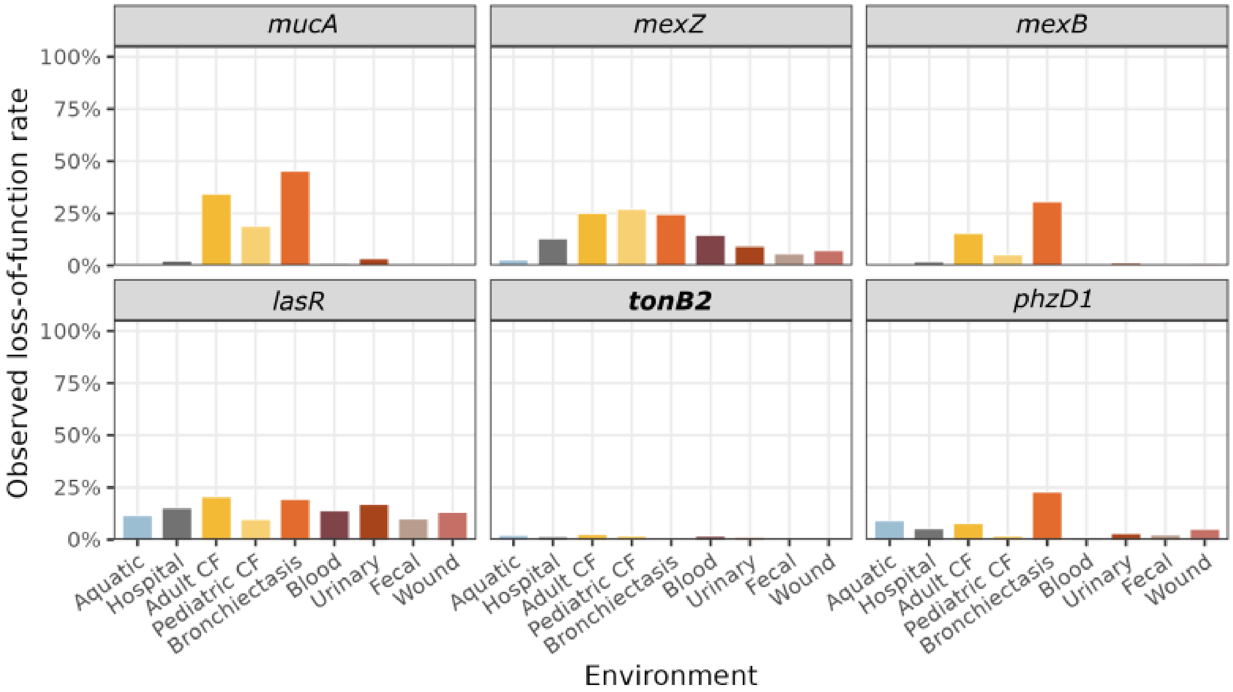


**Supplementary Fig. 8 | Relative abundance of loss-of-function mutations per environment and gene.** Putative loss-of-function mutations are defined as deletions (yielding ≤40% coverage), frameshifts, or nonsense mutations. The relative proportion of these mutation types per environment is plotted across genes of interest identified in the model SHAP analysis. This confirms that the model captures biologically meaningful variation associated with distinct environments.

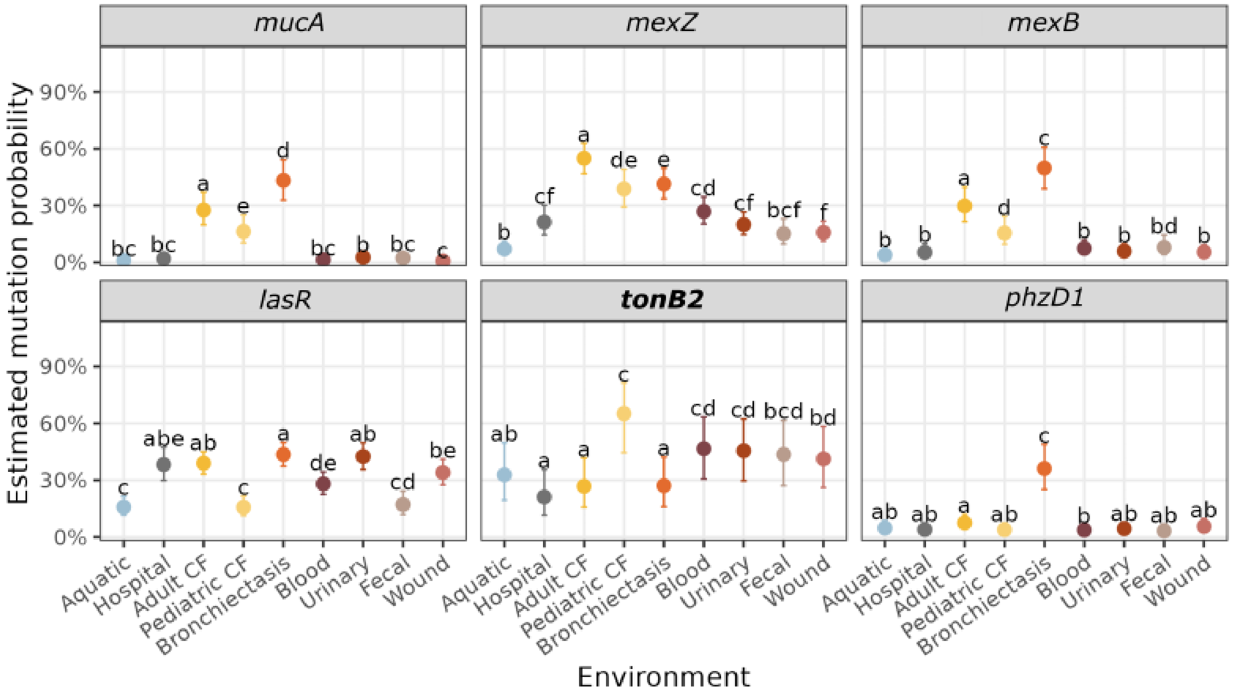


**Supplementary Fig. 9 | Mutation probability per gene excluding contig splits.** The genes tonB2 and phzD1 frequently exhibit contig splits, in which part of the gene spans the end of one contig and the beginning of another. While these may have biological explanations, we sought to ensure that putatively artifactual splits (for example, due to sequencing or assembly error) did not bias our results. For all top genes of interest, mutation probabilities were estimated from an LMER mixed-effects model (points ± SE) by environment across all P. aeruginosa genomes relative to the PAO1 reference, with genes showing any contig splits excluded. The model includes environment as a fixed effect and genomovar as a random effect, with Tukey post hoc tests used to assess pairwise differences.
